# A prion-like protein regulates the 2-dimensional to 3-dimensional growth transition in the moss *Physcomitrium patens*

**DOI:** 10.1101/2024.04.08.588603

**Authors:** Zoe Weeks, Gargi Chaturvedi, Emily Day, Steven Kelly, Laura A. Moody

**Affiliations:** Department of Biology, University of Oxford, South Parks Road, Oxford, OX1 3RB, United Kingdom

**Keywords:** 3D growth, plant development, apical cell, *Physcomitrium patens*

## Abstract

The colonization of the land by plants coincided with the evolution of 3-dimensional (3D) growth; the acquisition of apical cells with the capacity to rotate the plane of cell division. The moss *Physcomitrium patens* has recently been developed as a model to dissect the genetic basis of 3D growth, an invariable and unifying feature of all land plants. The cytokinin-unresponsive *Ppnog1-R* mutant incorrectly orients division planes in developing buds and thus fails to make the transition to 3D growth. To reveal the genetic interactors of the *PpNOG1* gene, which encodes a protein with a C-terminal UBA domain, we performed a screen and identified the *suppressor of nog1a* (*snog1a*) mutant. We have mapped the causative mutation to a gene that encodes a prion-like protein related to FLOE2/3 and demonstrated that the mutant phenotypes observed in both a *nog1* disruptant mutant (*nog1dis*) and *snog1a* can be attributed to changes in cytokinin perception. We present a revised model for 3D growth and suggest that the 2D-to-3D growth transition is regulated, at least in part, by liquid-liquid phase separation (LLPS).

**SUMMARY STATEMENT:** The transition to 3D growth is negatively regulated by a prion-like protein that both alters cytokinin signaling and has been implicated in liquid-liquid phase separation (LLPS).

## INTRODUCTION

In the absence of cell movement, plants rely on cell growth processes, combined with asymmetric and precisely orientated cell divisions, to generate new morphologies and diverse cell types with specialised functions. The main driving force behind cellular diversity is the formation of apical cells that can divide to self-renew and generate new cell types, and it is the geometry of an apical cell and the way it divides that can greatly influence the pattern of growth and development that follows. Thus, diversification of plant form can largely be attributed to altered division processes in apical cells, which occurred prior to the transition from water to land approximately 470 million years ago (Kenrick & Crane, 1997).

Charophycean green algae, the sister lineage to the land plants, can develop apical cells but these only have the capacity for either 1-dimensional (1D) growth (one cutting face) or 2-dimensional (2D) growth (two cutting faces). As a result, the morphologies represented within the multicellular charophytes are typically filamentous (e.g., *Chara braunii*) or disc-like (e.g., *Coleochaete orbicularis*) (Kenrick and Crane, 1997; Wickett et al., 2014; Delwiche and Cooper, 2015; Harrison, 2017). 3-dimensional (3D) growth, the development of apical cells with three or more cutting faces, is a unifying and unvarying feature of all land plants. It was likely the emergence of 3D growth processes, along with the development of a multicellular sporophyte and the acquisition of vegetative desiccation tolerance, that enabled successful terrestrialization (Delwiche & Cooper, 2015; Harrison, 2017; Moody, 2020).

The gametophyte of the moss *Physcomitrium patens* is well suited to studies of 3D growth. This is because an extensive 2D filamentous growth phase precedes the transition to 3D growth, and thus the 2D to 3D growth transition can be studied without causing lethality, as the 2D growth phase can be vegetatively propagated in perpetuity (Moody et al., 2018; Moody et al., 2018b; Moody et al., 2021). The gametophyte phase of the *P. patens* life cycle begins with the germination of a haploid spore, which gives rise to a 2D branching filamentous network known as the protonema, which extends by tip growth. The protonema consists of two cell types; the first to emerge are the chloroplast-dense chloronemal cells, and subsequently caulonemal cells that form because of an auxin-mediated reprogramming of a chloronemal apical cell into a caulonemal apical cell (Jang and Dolan, 2011; Jaeger and Moody, 2021). Caulonemal cells can undergo a lateral division to produce side branch initials, most of which go on to form secondary protonema (2D fate, approximately 95%) but some acquire 3D fate (gametophore initials) and give rise to gametophores (approximately 5%) (Aoyama et al., 2012). The fate of a side branch initial is likely determined by highly localized cues within a caulonemal cell prior to side branch emergence, as a single caulonemal cell can simultaneously divide to give rise to both a filament and a gametophore (Harrison et al., 2009). Elegant work by Aoyama and colleagues has demonstrated that persistent expression of a group of AP2-type transcription factors (*PpAPB1-4*) is required to commit a side branch to 3D cell fate. Loss of *PpAPB1-4* expression corresponds to the maintenance of 2D cell fate, and consequently, mutants lacking all four genes fail to make the transition from 2D to 3D growth (Aoyama et al., 2012).

A filament initial is readily distinguishable from a gametophore initial: (i) the angle of the lateral “transition” division can dictate whether a caulonemal cell gives rise to a filament or a gametophore (Tang et al., 2020); and (ii) gametophore initials swell diffusely and divide in a characteristically oblique manner during the specification of 3D growth, at the first, second and third divisions of the developing gametophore. Successive rotating divisions then lead to the establishment of a tetrahedral apical cell at the apex of the shoot. The apical cell continuously self-renews and divides in three planes to form phyllid initials that go on to form the leaf-like phyllids that are organized around the central axes of developing gametophores in a spiral phyllotaxy (Harrison et al., 2009). Mature gametophores bear both archegonia and antheridia, which house eggs and sperm respectively. Following fertilization, a mitotic programme initiates, leading to the formation of a multicellular diploid sporophyte, which undergoes meiosis to produce haploid spores to restart the life cycle (Cove and Knight, 1993).

It has long been known that cytokinin induces the formation of gametophore initials but is insufficient to maintain 3D growth, as treatment with high levels of cytokinin produces buds that develop into callus-like tissue, rather than structurally organized gametophores (Brandes and Kende, 1968; Ashton et al., 1979). In a variety of developmental contexts, auxin has been consistently implicated as the signal required to ‘break symmetry’ in plants (Petricka et al., 2009; Shao and Dong, 2016). Perhaps unsurprisingly, the formation of gametophore initial cells and the specification and maintenance of 3D growth also relies upon auxin signaling, although the application of high levels of auxin can antagonize cytokinin and reverse the effects of cytokinin treatment (Brandes and Kende, 1968). Nevertheless, auxin levels are notably high during the specification of 3D growth but diminish once a tetrahedral apical cell has been established (Thelander *et al*., 2018). It is likely that auxin acts to promote cell differentiation and that cytokinin acts to continually maintain apical cell proliferation (Hata and Kyozuka, 2021). Thus, it appears that a highly regulated balancing act between auxin and cytokinin is required for both the specification and maintenance of 3D growth.

In recent years, functional studies have demonstrated that the transition to 3D growth is complex and regulated at many levels; both epigenetically (Mosquna et al., 2009; Okano et al., 2009; Raquid et al., 2023) and transcriptionally (Aoyama et al., 2012), and an ever-expanding number of studies have begun to connect complex cell signaling pathways to post-translational regulation (Girod et al., 1999; Perroud et al., 2014; Demko et al., 2014; Johansen et al., 2016; Hoernstein et al., 2016; Schuessele et al., 2016; Whitewoods et al., 2018; Moody et al., 2018; Perroud et al., 2020; Moody et al., 2021; Cammarata et al., 2022). Using a forward genetics approach, we previously demonstrated that the *NO GAMETOPHORES 1* (*PpNOG1*) gene is essential for the transition to 3D growth in *P. patens*. Notably, the *PpNOG1* gene encodes a protein with a prominent C-terminal ubiquitin-associated (UBA) domain, which has been shown to be associated with protein degradation processes (Hofman and Bucher, 1996; Su and Lau, 2009; Moody et al., 2018). Mutants lacking a functional copy of *PpNOG1* (the ‘*no gametophores 1 – Reference’* mutant; *Ppnog1-R*) produce significantly fewer gametophore initials than wild type, even in the presence of cytokinin. Those gametophore initial cells that do form cannot correctly orient the characteristically oblique plane of the first cell division. Cell division planes are then misplaced thereafter, leading to the formation of defective gametophores that undergo very early developmental arrest (Moody et al., 2018). Since the *PpAPB* genes are downregulated in the *Ppnog1R* mutant, we previously proposed that PpNOG1 may positively regulate 3D growth by degrading a repressor of *PpAPB* transcriptional activation, although the identity of the target(s) remains unknown (Aoyama et al., 2012; Moody et al., 2018). To build on our understanding of the role played by *PpNOG1* in both the initiation and specification of 3D growth, we generated a *Ppnog1* disruption mutant (*Ppnog1dis*) and then performed a suppressor screen to identify mutations that alleviated the *Ppnog1dis* mutant phenotype (i.e., reversion to 3D growth). Here, we describe the screen, along with the detailed characterization of the *suppressor of nog1a* (*snog1a*) mutant and the identification of the causative mutation within a gene encoding a prion-like protein.

## RESULTS

### Disruption of the *PpNOG1* locus recapitulates the *Ppnog1-R* mutant phenotype

Previously, we performed a UV-mediated forward genetic screen that led to the identification of the *PpNOG1* gene (Moody et al., 2018). To explore the *PpNOG1* genetic interaction network underpinning the 2D to 3D growth transition, we designed a suppressor screen to identify mutations that alleviated the 3D-defective phenotype caused by loss of *PpNOG1* function (i.e., the reacquisition of 3D growth). A preliminary suppressor screen of approximately 3,000 UV-mutagenized lines of the original *Ppnog1-R* mutant, generated in our original forward genetic screen, identified eight mutants that exhibited a complete reversion to 3D growth. In each of these mutants, the phenotype was caused by correction of the previously described mutation; a T>C transition that converted the premature termination codon back to an arginine residue (data not shown). We therefore decided to disrupt the *PpNOG1* locus in such a way that would prevent repair simply through the introduction of UV-induced SNPs. To that end, we set out to generate a *nog1* knockout mutant in which the entire coding sequence of *PpNOG1* had been replaced with a hygromycin resistance cassette. Several attempts were made to obtain a line in which the entire *PpNOG1* sequence had been removed. However, as was the case in previous attempts (Moody et al., 2018), we could achieve disruption of the *PpNOG1* locus but unfortunately not a full deletion. Nevertheless, a *nog1* null mutant was generated in which a portion of the *PpNOG1* promoter had been excised following recombination (Fig. S1A). Furthermore, we were consistently unable to detect a *PpNOG1* transcript in this line, and thus had confirmed that we had generated a bona fide *PpNOG1* disruptant mutant (*nog1dis*) (Fig. S1B). Consistent with the original *Ppnog1-R* mutant, the *nog1dis* mutant completely and consistently failed to make the transition to 3D growth (Fig. 1). Thus, disruption of the *PpNOG1* locus recapitulated the original *Ppnog1-R* mutant phenotype and generated a suitable line for mutagenesis.

**Figure 1.**
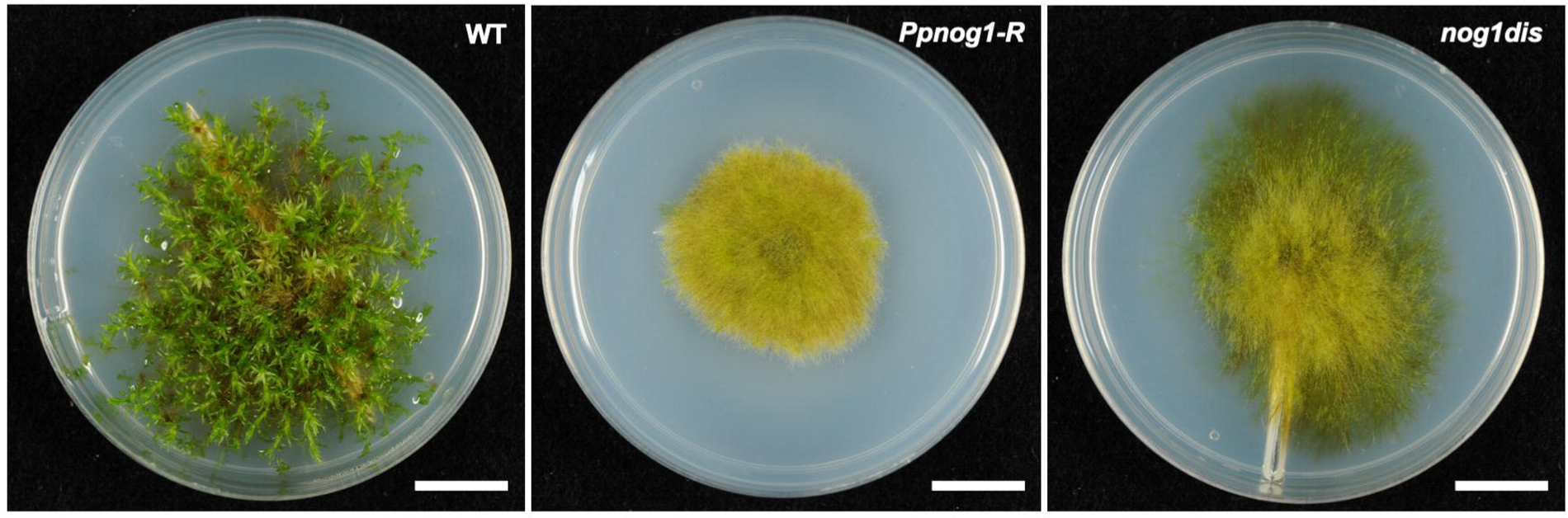
The nog1dis mutant fails to make the transition to 3D growth. Representative images of 6-week-old Villersexel wild type (WT), *Ppnog1-R* and *nog1dis* plants showing the presence (WT) and absence (*Ppnog1-R* and *nog1dis*) of gametophores. Scale bars, 1 cm.

### The *suppressor of nog1a* (*snog1a*) mutant can specify 3D growth and responds to cytokinin

A suppressor screen of 2,864 UV-mutagenized lines of the *nog1dis* mutant yielded two ‘*suppressor of nog1*’ (*snog1*) mutants that exhibited a restoration of 3D growth. In one of these mutants, *suppressor of nog1a* (*snog1a*), the formation of gametophores was partially restored to approximately 45% of the frequency of wild type (Fig. 2A,B), although these were somewhat stunted and emerged later than those formed in the wild type (Fig. 2C,D; Fig. S2).

**Figure 2.**
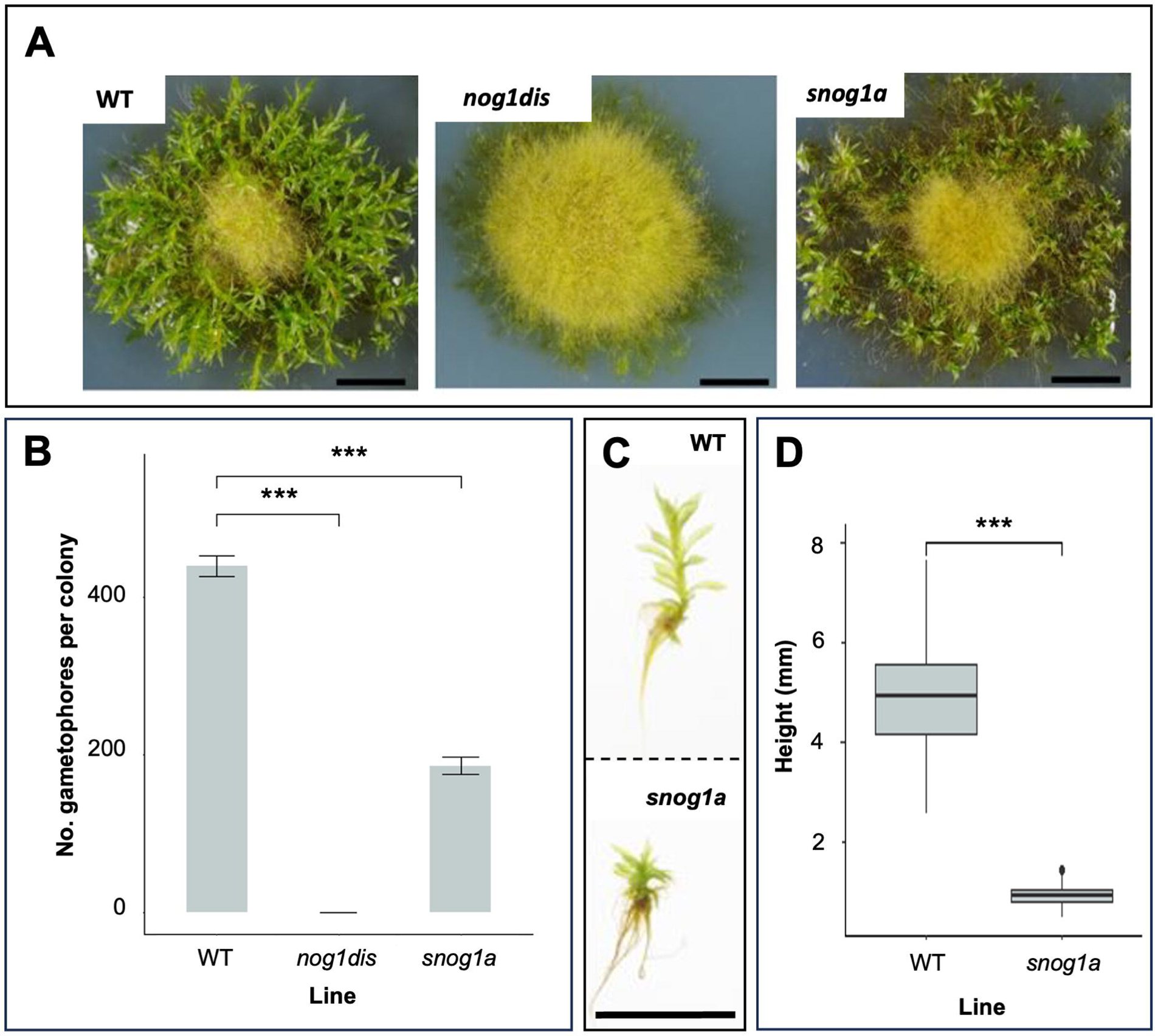
The *snog1a* mutant exhibits a partial restoration of 3D growth. A) Representative images of 6-week-old Villersexel wild type (WT), *nog1dis* and *snog1a* plants showing the presence (WT and snog1a) and absence (nog1dis) of gametophores. B) Mean number of gametophores per culture (n=5) ± SEM (t test ***p < 0.05). C) Representative images of a gametophore from wild type (top) and stunted gametophore from the *snog1a* mutant (bottom). D) Mean height of gametophores from wild type (n=100) and *snog1a* (n=80) ± SEM (t test ***p < 0.05). Scale bars, 1 cm (A and C).

In contrast to the *nog1dis* mutant, in which responses to cytokinin are impaired in a similar manner to that of the *Ppnog1-R* mutant, buds can be induced by cytokinin treatment in the *snog1a* mutant, although to a lesser extent than in wild type (Fig. 3). Thus, the mutant phenotypes observed in both the *nog1dis* and *snog1a* mutants can be attributed to changes in cytokinin perception.

**Figure 3.**
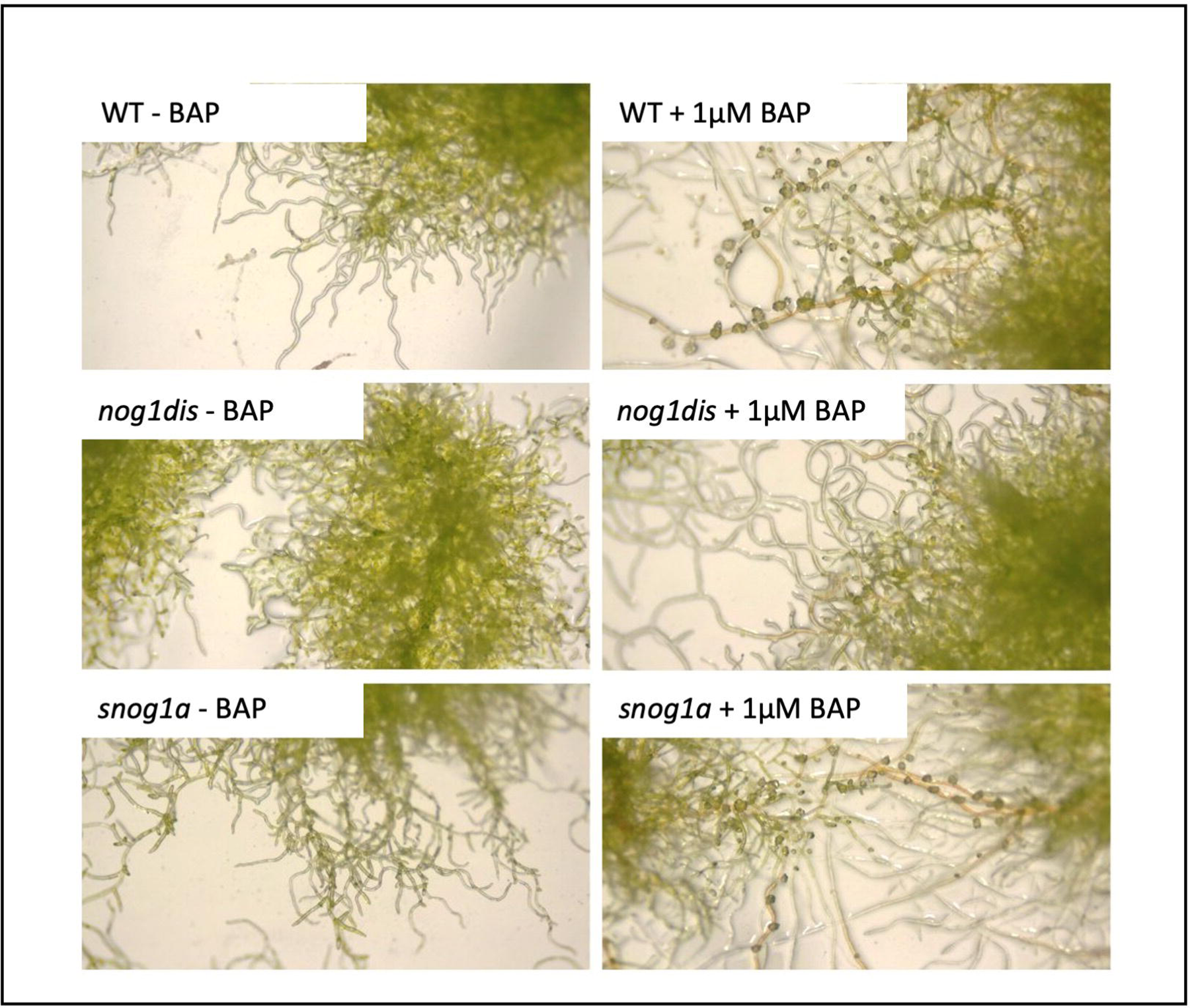
The *snog1a* mutant is cytokinin responsive. Representative images of wild type (WT), *nog1dis* and *snog1a* plants cultured in the presence or absence of the cytokinin analogue 6-benzylaminopurine (BAP).

To determine whether the cell division orientation defects had been repaired in the *snog1a* mutant, we obtained z-stack projections of developing buds stained with propidium iodide. In wild-type *P. patens*, the first division of the gametophore initial cell was invariably oblique and yielded an apical and a basal cell (Fig. 3A). Two additional oblique divisions of the apical and basal cells then occurred rather synchronously. Although the angle of the division plane was consistent in each case, the order in which the cells divided was inconsistent; we generally observed that the apical cell divided before the basal cell as often as the basal cell divided before the apical cell (Fig. 3B,C). Successive rotating divisions specified a gametophore apical cell with a characteristic tetrahedral shape, which self-renewed and divided to give rise to the phyllids, which wrap around the central axis of the gametophore in a spiral phyllotaxy (Fig. 3D) (Harrison et al., 2009). In the *nog1dis* mutant, similarly to the previously described *Ppnog1-R* mutant, significantly fewer gametophore initial cells formed, and the first division plane of the gametophore initial cell was not characteristically oblique. In most cases the initial division pattern followed that of a filament initial cell, in which the division plane was positioned roughly parallel to the parental cell from which it was derived (Fig. 3E). Cell plates were then positioned randomly during subsequent divisions, which prevented the specification and maintenance of a tetrahedral apical cell. In some cases, the gametophore initial cells swelled and elongated excessively but could then divide in a somewhat oblique manner at the first and second divisions (Fig. 3F). However, cell plates were invariably misplaced from the onset of the third division (Fig. 3G). Most developing gametophores arrested early in development, but occasionally callus-like buds appeared because of uncontrolled proliferation (Fig. 3H). Moreover, bifurcation events were occasionally observed, which were a result of confused cell fate acquisition, and often associated with supernumerary apical cell formation. However, none of these apical cells were successfully maintained long enough to produce a gametophore. In the *snog1a* mutant, the formation of gametophore initial cells was partially restored, and these largely followed a wild-type pattern of development, with some exceptions. In some cases, we observed that the angle of the first division plane had been corrected to some extent, but not fully, and in these instances a mature gametophore was not formed (Fig. 3I). However, in most cases, the first division plane was characteristically oblique and thus the orientation of the division plane had been fully corrected (Fig. 3J). This was followed by two further correctly oriented oblique divisions (Fig. 3K), which formed the prerequisites for the specification of a conspicuous tetrahedral apical cell at the apex of developing gametophores (Fig. 3L). Thus, restoration of gametophore initial cell formation in the *snog1a* mutant was accompanied by the reversion of the cell division orientation defects observed in the *nog1dis* mutant.

Notably, the phenotype observed in the *snog1a* mutant only constituted a partial restoration of the *nog1dis* phenotype. Nevertheless, a tetrahedral apical cell was established and maintained in the *snog1a* mutant, and the gametophores formed were fully viable.

### The causative mutation of *snog1a* resides in a gene that encodes a prion-like protein

To identify the causative mutation in the reproductively viable *snog1a* mutant, we obtained phenotypically segregating populations by performing a cross between the *snog1a* mutant and the highly fertile non-mutagenized Reute::mCherry strain (Perroud et al., 2020). Because a conventional cross between two different haploid strains was carried out, and at least two genetic loci were mutated (the mutation within the *PpNOG1* gene, and the unknown mutation that caused the *snog1a* phenotype), we expected one quarter of the progeny to exhibit the original *nog1dis* mutant phenotype and the remainder to exhibit varying capacities for 3D growth (Fig. S3A). The frequencies that we observed were generally consistent with the mutation of a single genetic locus in the *snog1a* mutant, although this is not statistically significant (Fig. 5A). One possible explanation for the reduced *nog1dis* mutant phenotype frequency observed was that disruption of the *PpNOG1* gene affects haploid spore germination and/or the viability of young sporelings. Nevertheless, the frequencies observed did not support the idea that causative mutations resided in two or more genetic loci. We therefore prepared genomic DNA from 80 individuals that exhibited the *nog1dis* phenotype and pooled these in equimolar amounts, and this became the ‘wild-type pool’ (all individuals resembled the non-mutagenized parental line, *nog1dis*). We also prepared genomic DNA from 98 individuals with the capacity for 3D growth, showing a preference for those that most closely resembled the *snog1a* mutant, and this became the ‘mutant pool’ (Supplementary Figure S3A). These two pools were sequenced at 44X coverage alongside both parental lines; the *nog1dis* mutant (generated in the Villersexel wild-type strain) and the Reute::mCherry line. To identify the genomic region containing the *snog1a* mutation, bulk segregant analysis was performed. Single nucleotide polymorphisms (SNPs) that differed between the two parental lines were identified as markers and the frequency of each SNP variant was mapped across the chromosomes to show the parental origin of each region. For regions not associated with the phenotypes in the two pools, the expected SNP frequency was around 0.5, showing equal contribution from both parents. The expected *snog1a* mutant allele frequency was 0 in the wild-type pool and 0.67 in the mutant pool (Fig. S3B,C). When the allele frequencies for the mutant individuals were plotted across all 27 chromosomes in the *P. patens* genome assembly, a peak of the expected allele frequency was revealed on chromosome 8, which provided a region of the genome to interrogate for causative mutations (Fig. S3D). Our analysis revealed that two C > T transitions generated two distinct in-frame termination codons (Gln^335^Ter and Gln^374^Ter) in a single gene (Pp3c8_19720) (Fig. 5B,C). We cloned and sequenced the coding sequence of this gene and confirmed that both point mutations were present in the *snog1a* mutant, but absent from both the *nog1dis* mutant and the Reute::mCherry line (Supplementary Data S1). In addition, sequencing of the corresponding transcript confirmed the presence of the point mutation but did not reveal any splice variants in the *snog1a* mutant, a phenomenon observed in both previously described *Ppnog1-R* and *Ppnog2-R* mutants (Supplementary Data S2) (Moody et al., 2018; Moody et al., 2021).

**Figure 4.**
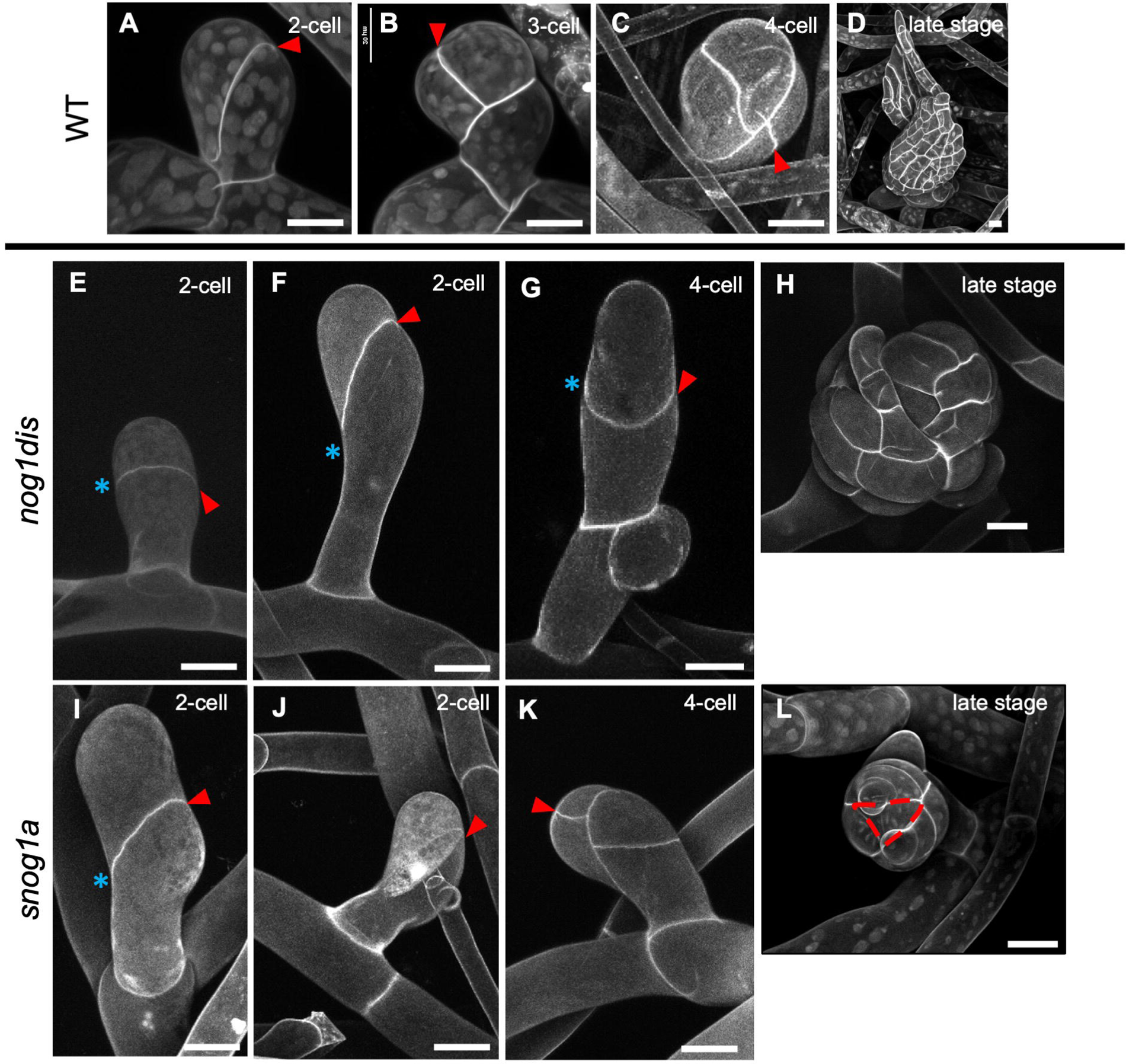
The *snog1a* mutant can establish and maintain a tetrahedral apical cell. Propidium-iodide-stained buds of wild type at the 2-cell (A), 3-cell (B) and 4-cell (C) and late stage (D); the nog1dis mutant at the 2-cell (E,F), 4-cell (G) and late stage (H); and the snog1a mutant at the 2-cell (I,J), 4-cell (K) and late stage (L). Red arrows denote the most recent division in each developing bud, and blue asterisks highlight misoriented division planes in *nog1dis* or *snog1a* mutants.

**Figure 5.**
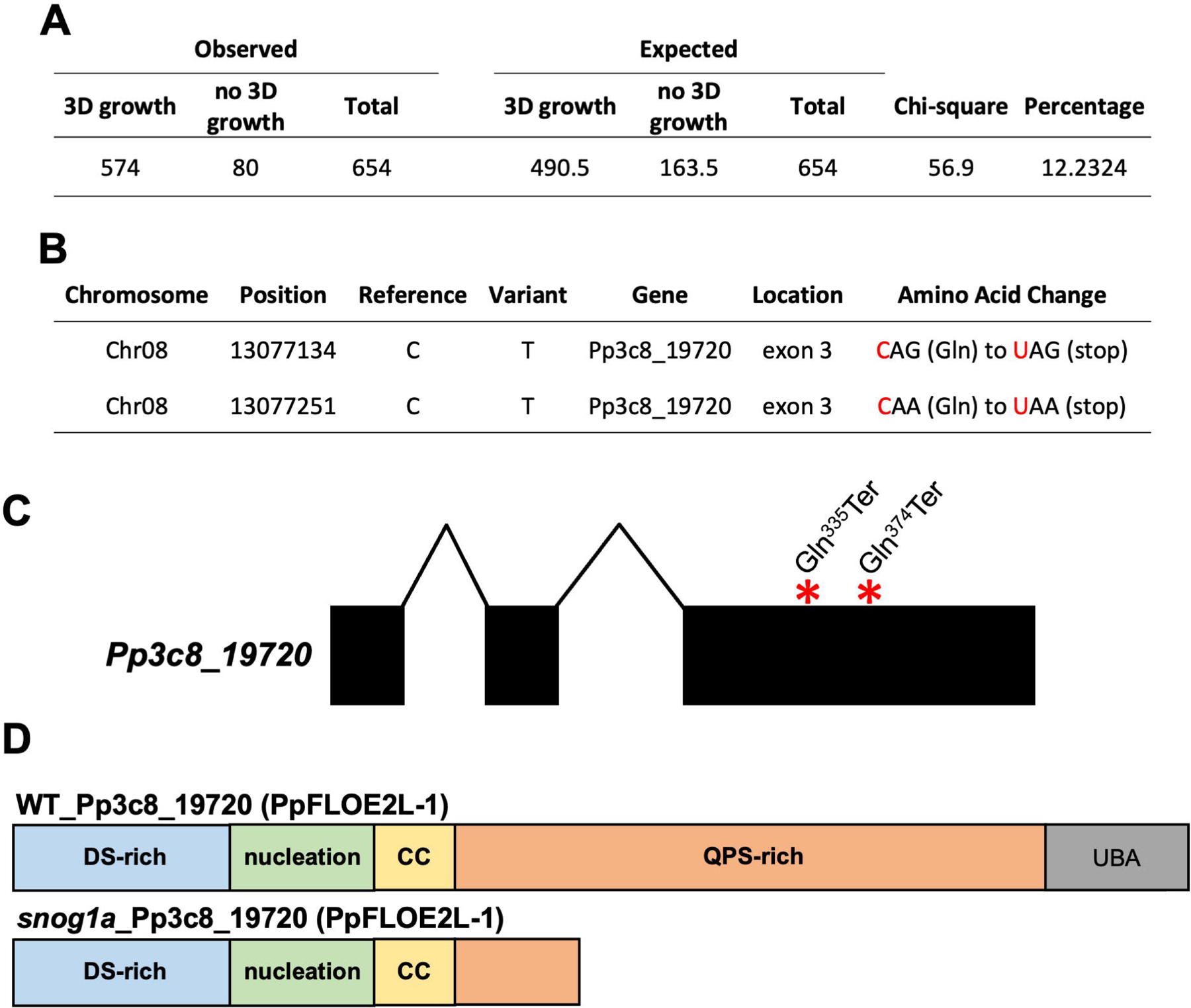
Identification of the causative mutation in the *snog1a* mutant. A) Phenotypic analysis of spore progeny derived from a cross between *snog1a* and the Reute::mCherry wild-type strain. B) Gene candidates identified following interrogation of the genomic locus on chromosome 8. C) Gene structure diagram of *Pp3c8_19720* highlighting the presence of two termination codons in the third exon (exons – blocks, introns – horizontal lines). D) The wild-type Pp3c8_19720 protein (top) and the truncated Pp3c8_19720 protein in *snog1a* (bottom).

The gene mutated in the *snog1a* mutant encodes a protein with a C-terminal UBA, and thus the protein domain architecture resembles that of the PpNOG1 protein (Moody et al., 2018). To infer phylogenetic relationships for the putative SUPPRESSOR OF NOG1A protein, we set out to retrieve orthologous sequences from the genomes of representatives of the chlorophytes, charophytes, bryophytes, lycophytes, monilophytes, gymnosperms and angiosperms (Supplementary Data S3). Remarkably, Pp3c8_19720 shares homology with an Arabidopsis prion-like protein (AtFLOE1) that can undergo hydration-dependent phase separation (PS) to regulate seed germination, and related proteins (AtFLOE2 and AtFLOE3) that have been shown to undergo PS in a transient expression system (Dorone et al., 2021). As with the analyses conducted by Dorone *et al*. (2021), our own phylogenetic analyses, revealed the presence of two discrete clades, a FLOE1-like (FLOE1L) and a FLOE2-like (FLOE2L) clade. However, while it was previously stated that the FLOE1L clade was restricted to seed plants, we found FLOE1L representatives in the seed plants in addition to the monilophytes. Conversely, the FLOE2L clade included Pp3c8_19720, as well as three additional *P. patens* paralogues, and homologues in most land plants except for hornworts, where they appear to have been lost. FLOE2L homologues were also identified in some green algal lineages (Fig. S4, Supplementary Data S3). Given the significant homology between Pp3c8_19720 and representatives within the FLOE2L clade, we hereafter refer to Pp3c8_19720 as PpFLOE2L-1.

The AtFLOE1, AtFLOE2 and AtFLOE3 proteins contain a predicted folded domain (known as the nucleation domain), a coiled-coil domain (CC), a UBA domain at the C-terminus and two disordered regions; one enriched for aspartic acid and serine (DS-rich domain.) and one enriched for glutamine, proline, and serine (QPS-rich domain) (Dorone et al., 2021). Alignment of PpFLOE2L-1 with AtFLOE1, 2 and 3 confirmed that these protein domains were conserved in PpFLOE2L-1 (Fig. 5D, Fig. S5). As a result of the nonsense mutations introduced into *PpFLOE2L-1*, a truncated protein was formed in *snog1a*, which was missing most of the QPS-rich domain in addition to the UBA (Fig. 5D; Supplementary Data S2).

To confirm that we had correctly identified the causative mutation, we used the rice actin promoter to drive the expression of a wild-type version of *PpFLOE2L-1* in the *snog1a* mutant. The resulting line exhibited a reversion to the 3D-defective phenotype seen in the *nog1dis* mutant (Fig. 6A,B and Fig. S6). We also generated two independent lines in which *PpFLOE2L-1* was deleted in the *nog1dis* mutant. Each line exhibited a restoration of 3D growth, and thus the *snog1a* mutant phenotype had successfully been recapitulated (Fig. 6C-E, Fig. S7). Thus, the causative mutation of *snog1a* resides in a gene that encodes a prion-like protein.

**Figure 6.**
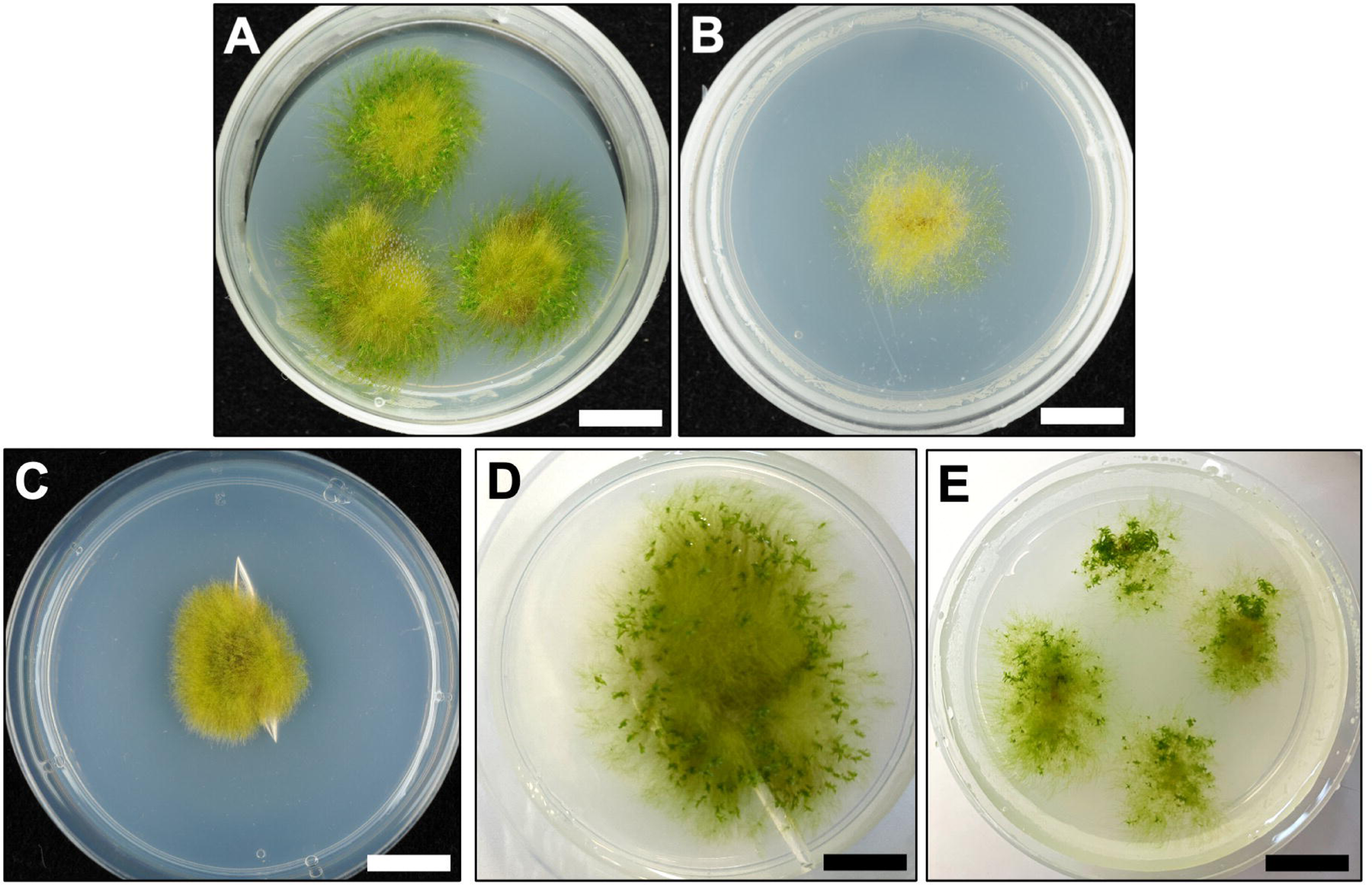
Confirmation of the causative mutation in the *snog1a* mutant. (A,B) Representative images of 6-week-old snog1a (A) and snog1a complemented with pAct::PpFLOE2L-1 (B). (C-E) Representative images of 6-week-old nog1dis (C) and nog1dis/floe2l-1_4 (D) and nog1dis/floe2l-1_6 (E) double disruptants. Scale bars, 1 cm.

## DISCUSSION

The acquisition of apical cells with the capacity for 3D growth occurred in the last common ancestor of land plants, and enabled the diverse morphologies seen across the planet today (Kenrick and Crane, 1997; Delwiche and Cooper, 2015). We previously showed that the *NO GAMETOPHORES 1* (*PpNOG1*) gene is required for 3D growth in *P. patens*, an extant representative of the bryophytes. Notably, in mutants lacking *PpNOG1* function (*Ppnog1-R*), gametophore initial cell formation is significantly reduced, even in the presence of cytokinin. Furthermore, cell division planes are misoriented in emerging gametophores, which subsequently undergo premature developmental arrest. Thus, *PpNOG1* promotes the cytokinin-mediated transition from 2D to 3D growth and positively regulates the orientation of cell divisions required to establish a tetrahedral apical cell (Moody et al., 2018).

To reveal new insights into the genetic interaction network underpinning the transition from 2D to 3D growth, we first recapitulated the *Ppnog1-R* mutant phenotype in an independent *Ppnog1* disruption mutant (*nog1dis*), in which *PpNOG1* was not expressed (Fig. 1). The *nog1dis* mutant was generated using a targeted approach, and thus lacked the background SNPs that were introduced into the *Ppnog1-R* mutant by UV-based mutagenesis. We then performed a UV-mediated suppressor screen to identify mutations that alleviated the *nog1dis* mutant phenotype to permit the reversion to 3D growth. The primary focus of this paper is the characterisation of the ‘*suppressor of nog1a’* (*snog1a*) mutant identified in this screen.

Similarly to the original *Ppnog1-R* mutant, the *nog1dis* mutant was unable to form gametophores even in the presence of cytokinin (Fig. 1,3). Mutants in which all four *PpAPB* genes have been disrupted exhibit a similar level of cytokinin-unresponsiveness to both *Ppnog1-R* and *nog1dis* mutants and fail to make the 3D growth transition (Aoyama et al., 2012). Furthermore, we previously observed that *PpAPB* genes are downregulated when *PpNOG1* is absent (Moody et al., 2018). Thus, *PpNOG1*, along with the *PpAPB* genes are integral regulators of cytokinin perception during the switch from 2D to 3D growth. Since the *snog1a* mutant exhibits responsiveness to cytokinin, this suggests that those cytokinin signaling components, ordinarily repressed in *nog1dis*, have been reactivated in the *snog1a* mutant. This demonstrates that the phenotypes observed in both *nog1dis* and *snog1a* can be attributed to alterations in cytokinin perception. Thus, this study has begun to shed light on the potential mechanism underlying cytokinin-mediated 3D growth initiation, a topic that has fascinated biologists for several decades (Brandes and Kende, 1968; Ashton et al., 1979; Reski and Abel, 1985; Schulz et al., 2000; Schulz et al., 2001; von Schwartzenberg et al., 2007; von Schwartzenberg et al., 2016; Cammarata et al., 2022).

Although the formation of gametophore initial cells was not fully restored in the *snog1a* mutant, those that formed usually followed a wild-type pattern of development to achieve the establishment of a tetrahedral apical cell (Fig. 4). It has previously been shown that the positioning of cell division planes in emerging gametophores is dependent on microtubules (Kosetsu et al., 2017; Kozgunova et al., 2022). Thus, it is likely that the cell division orientation defects observed in the *nog1dis* mutant are due to aberrant organisation of the microtubule cytoskeleton. Since the correction of division plane orientation observed in the *snog1a* mutant is accompanied by the restoration of a cytokinin response, we speculate that that microtubule organisation that occurs during the 3D growth transition is dependent on cytokinin. This is consistent with other reports in the literature (Montesinos et al., 2020).

There is an increasing volume of literature that describes the regulatory role of cuticle-related genes in the 3D growth transition (Renault et al., 2017; Lee et al., 2020; Kreigshauser et al., 2021; Moody et al., 2021). These mutants invariably exhibit perturbations in the frequencies of gametophore initial cells that form, and those that do form exhibit division orientation defects that arrest gametophore development. Furthermore, the expression of these genes has been shown to be induced by cytokinin (Moody et al., 2021). Although we understand that a cuticle is absent in the protonema but observed in gametophores, the developmental stage at which cuticle biosynthesis initiates remains unclear (Renault et al., 2017; Lee et al., 2020; Kreigshauser et al., 2021). In several of the 3D-defective mutants described in the literature so far, including the *Ppnog1-R* mutant, the cuticle is invariably absent (Aoyama et al., 2012; Renault et al., 2017; Moody et al., 2018; Kreigshauser et al., 2021; Zhang et al., 2024). Thus, it is possible that the acquisition of the cuticle was a pre-requisite for 3D growth, and that the mechanical constraints imposed by the cuticle alter the division properties of cells formed in the steps toward the establishment of a tetrahedral apical cell. To support this hypothesis, the cuticle is notably absent in the *nog1dis* mutant but has been partially restored in the *snog1a* mutant, in which the partial reversion to 3D growth has occurred.

Our mapping approach and phylogenetic analyses revealed that the gene mutated in the *snog1a* mutant (Pp3c8_19720) was a homologue of the previously characterized FLOE genes in Arabidopsis; AtFLOE1, AtFLOE2 and AtFLOE3 (Fig. S4) (Dorone et al., 2021). Dorone et al. have demonstrated that AtFLOE1 undergoes reversible hydration-dependent liquid-liquid phase separation (LLPS) in the embryo, and that this process is dependent on the presence of a QPS-rich disordered domain. LLPS drives the formation of AtFLOE1 condensates when a seed is hydrated, but AtFLOE1 remains dispersed in desiccated seeds. Since loss of AtFLOE1 function can promote germination during drought or salt stress, it is thought that AtFLOE1 functions to inhibit seed germination in unfavourable conditions. The authors also demonstrated that AtFLOE2 and AtFLOE3, in addition to representatives from algae and bryophytes, undergo LLPS in a transient expression system (Dorone et al., 2021).

AtFLOE1 is a member of the FLOE1-like (FLOE1L) clade, which contains only those representatives from the monilophytes and seed plants. On the other hand, AtFLOE2 and AtFLOE3 reside within the FLOE2-like (FLOE2L) clade, which contains representatives from algal species as well as all land plant lineages. This includes Pp3c8_19720 (PpFLOE2L-1), in addition to three additional homologues in *P. patens*. We were able to complement the *snog1a* mutant phenotype with a full-length version of the wild-type coding sequence and recapitulate the phenotype by generating two independent double disruptant mutants (Fig. 6). The confirmation of gene identity enabled us to amend the name of *Pp3c8_19720* to PpFLOE2L-1. The presence of three additional *FLOE2L* homologues in the *P. patens* genome suggests that these genes may function redundantly. Thus, we hypothesise that higher order mutants of these genes will exhibit a progressive restoration of 3D growth, and indeed cuticle formation. FLOE2L genes were notably absent from hornworts, which suggests that the genetic toolkit underpinning 3D growth processes in hornworts is somewhat distinct.

Liquid-liquid phase separation (LLPS) is a phenomenon that has been increasingly linked to important developmental processes in plants (Fang et al., 2019; Huang et al., 2021; Cao et al., 2023). Furthermore, there have been reports that LLPS enables ubiquitin-binding shuttle proteins (i.e., those containing UBAs) to degrade ubiquitinated substrates (Dao and Castaneda, 2020). Similarly, to the three other *FLOE2-like* genes identified in *P. patens*, *PpFLOE2L-1* encodes a protein that contains a disordered domain enriched in glutamine, proline, and serine (QPS-rich), previously shown to serve as a prerequisite for LLPS; coiled-coil (CC) and nucleation domains; a disordered domain enriched in aspartic acid and serine (DS-rich); and a UBA. The presence of the UBA suggests that PpFLOE2L-1 plays a role in protein degradation, which is curiously reminiscent of the PpNOG1 protein. Due to the presence of two in-frame premature stop codons in the *PpFLOE2L-1* transcript, the *snog1a* mutant lacks most of the QPS-rich disordered domain as well as the UBA (Fig. 5). Thus, we hypothesise that PpFLOE2L-1 undergoes cell-type dependent LLPS to compartmentalize the cellular components required to degrade a ubiquitinated repressor of the 2D to 3D growth transition. In support of this notion, it has been reported that E3 ligases involved in protein degradation processes are able to function within condensates. Thus, we propose LLPS as a mechanism by which the induction of 3D growth can be rapidly triggered in response to both intrinsic and extrinsic cues, at the correct stage of development.

## SUMMARY

Since the loss of *PpFLOE2L-1* function can reverse the 3D-defective phenotype of the *nog1dis* mutant phenotype, we have demonstrated that *PpFLOE2L-1* acts as a negative regulator of 3D growth. Thus, we propose that PpNOG1 acts upstream of, and represses the activity of PpFLOE2L-1, which in turn induces the degradation of protein(s) that repress *PpAPB* gene transcription. We speculate that the decision to induce 2D versus 3D growth is dependent on the biophysical state of PpFLOE2L-1, and that these changes dynamically alter in a spatial and temporal manner; condensed when a side branch acquires 2D fate and dispersed when a side branch acquires 3D fate. We also propose that the transcriptional targets of the PpAPBs trigger the cytokinin response required to initiate the 3D growth transition. In the presence of cytokinin, cuticle-related genes are induced, which we propose subsequently triggers the induction of a CLAVATA-dependent auxin response to diminish the cytokinin response and prevent the inappropriate induction of 3D growth (Fig. 7).

**Figure 7.**
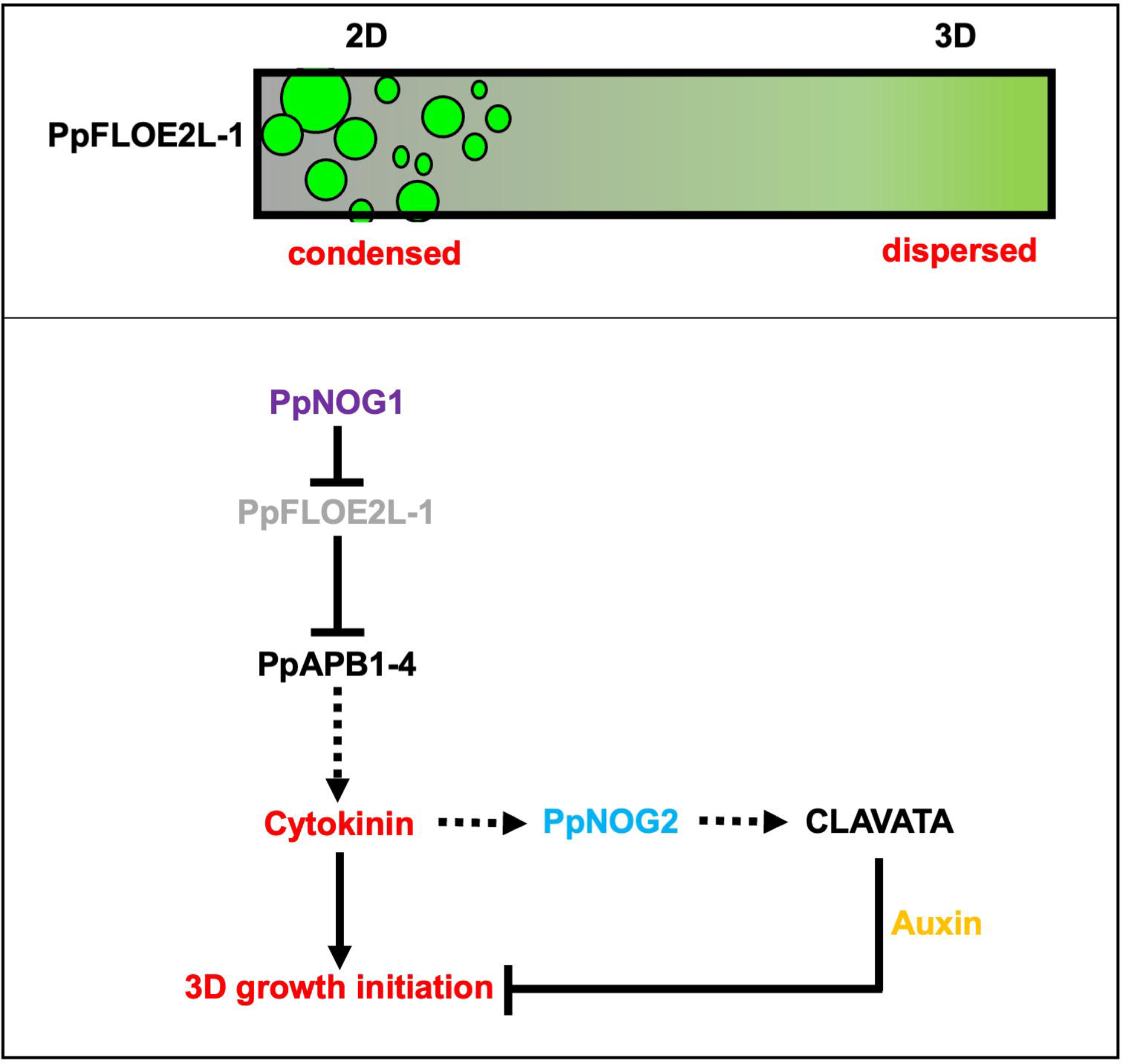
Speculative model for 3D growth regulation in *P. patens*. Top panel shows possible mechanism for cell-type specific LLPS of PpFLOE2L-1. Bottom panel shows possible relationship between PpFLOE2L-1 and other known regulators of 3D growth. PpNOG1 is proposed to act upstream of, and negatively regulate PpFLOE2-1, which in turn negatively regulates the expression of the *PpAPB* genes. Activation of the cytokinin signaling triggers the expression of cuticle-related genes, which act upstream of CLAVATA signaling components, which trigger an auxin-mediated repression of ectopic bud formation.

## MATERIALS AND METHODS

### Physcomitrium patens growth conditions

To encourage bud development, *P. patens* was grown on cellophane-overlaid BCD medium (1 mM MgSO_4,_ 1.84 mM KH_2_PO_4_ (pH 6.5), 10 mM KNO_3_, 45 µM FeSO_4_; 1 mM CaCl_2_; 0.1% Trace Elements Solution: 116 µM AlK(SO_4_)_2_, 220 µM CuSO_4_, 10 mM H_3_BO_4_, 235 µM KBr, 660 µM LiCl, 230 µM CoCl_2_, 190 µM ZnSO_4_, 2 mM MnCl_2_, 170 µM KI, 124 µM SnCl_2_) containing 0.8% agar. For routine propagation, and to stimulate filamentous growth, *P. patens* was grown on BCD medium supplemented with 5 mM ammonium tartrate (BCDAT). To enable propagation, tissues were harvested and homogenized in sterile water using an IKA T-25 digital ULTRA-TURRAX®. Tissues were then pipetted onto cellophane-overlaid BCDAT plates in a laminar flow hood. Plates were then placed in a growth cabinet at 24°C with a 16 h:8 h light (300 µmol m^-2^ s^-1^): dark cycle. Protoplasts were regenerated on cellophane-overlaid Protoplast Regeneration Medium (PRMB; BCDAT supplemented with 10 mM CaCl_2_, 0.5% glucose and 6% (w/v) D-mannitol) (Cove et al., 2009).

### Generation of the *nog1dis* mutant

A genomic DNA fragment from -827bp upstream of the start codon, up to but excluding the start codon of the *PpNOG1* sequence, was PCR-amplified using NOG1.5FKpnI and NOG1.5RXhoI primers (Table 1) and ligated into KpnI/XhoI cut pAHG1 (a kind gift from Yasuko Kamisugi and Andrew Cuming) to create pAHG1-*NOG1*-5’. A *PpNOG1* genomic DNA fragment including the stop codon up to 1420bp downstream of the *PpNOG1* stop codon was PCR-amplified using NOG1.3FNotI and NOG1.3RNotI primers and ligated into NotI cut pAHG1-*NOG1*-5’ to create *pNOG1*delH. *pNOG1*delH was linearized using KpnI prior to transformation into protoplasts isolated from the Villersexel wild type strain. Stable transformants were selected using 15 mgmL^-1^ Hygromycin B (Sigma-Aldrich cat. no. H9773).

**Table 1.**
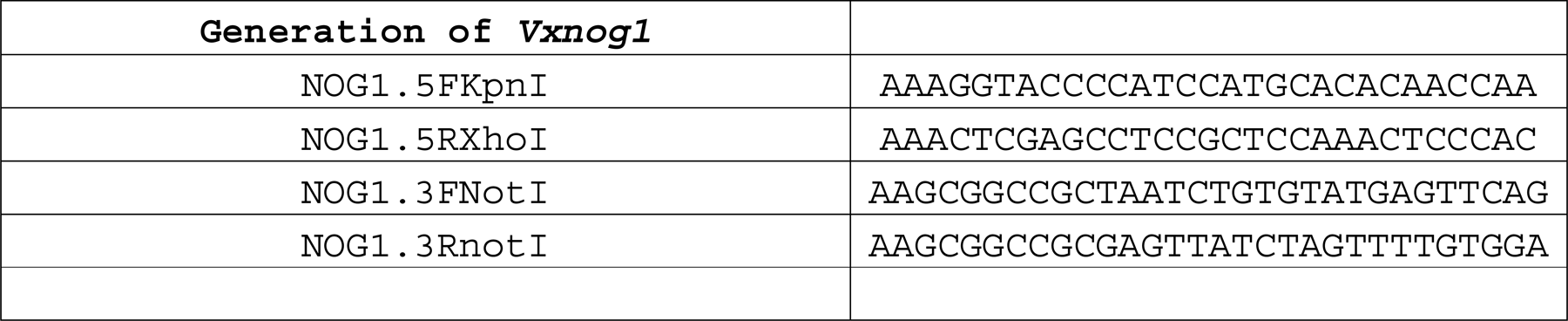
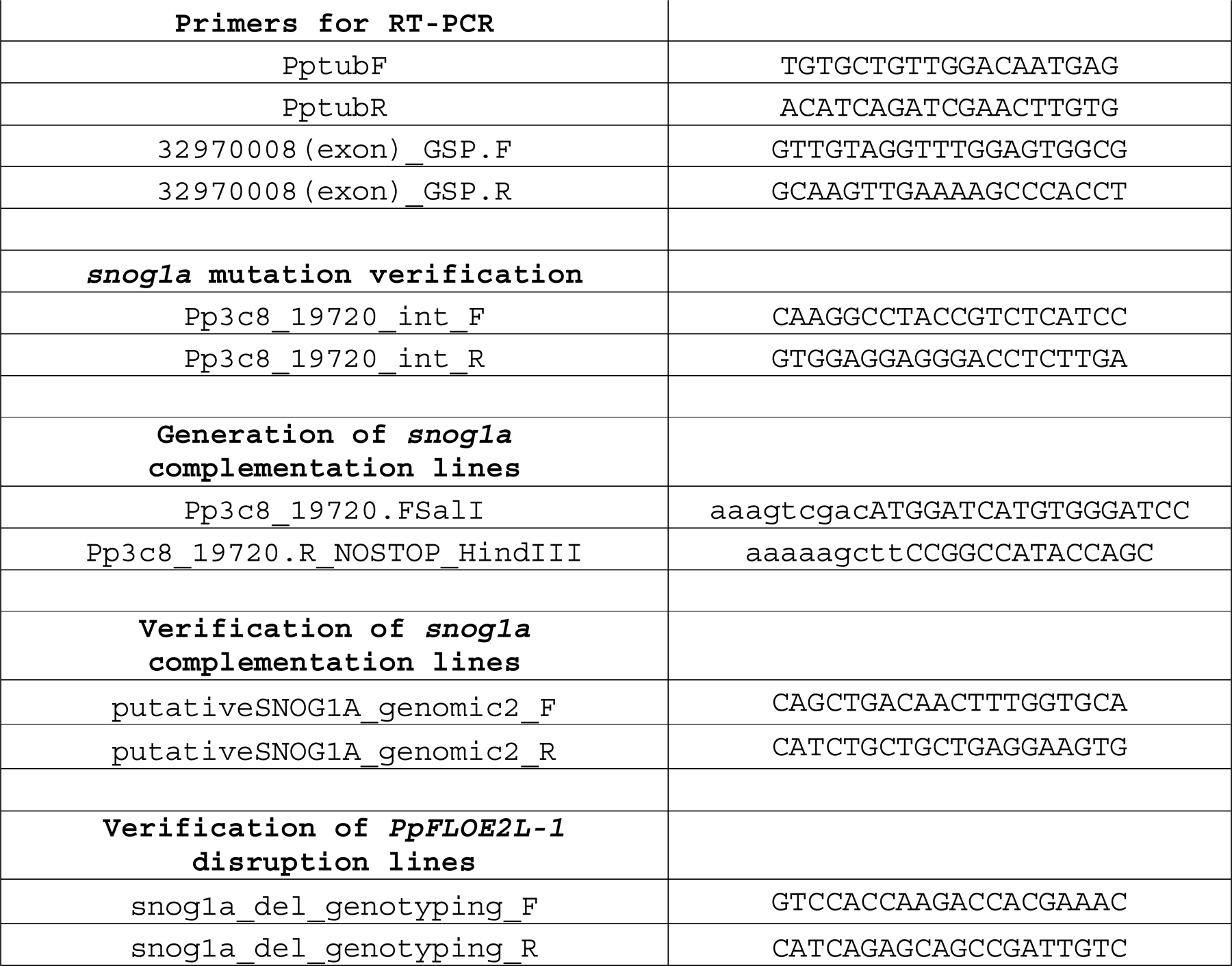
List of Primers.

### RT-PCR to detect absence of *NOG1* in the *nog1dis* mutant

Total RNA was isolated from two-week-old Villersexel wild type and the *nog1dis* mutant using the RNeasy kit (Qiagen) according to the manufacturer’s instructions. DNase treatment was carried out using TURBO DNase (Ambion) and cDNA synthesis performed using Superscript^TM^ III (ThermoFisher Scientific), as specified by the manufacturers. The *tubulin* transcript was amplified using PptubF and PptubR, and the *PpNOG1* transcript was amplified using 32970008(exon)_GSP.F and 32970008(exon)_GSP.R (Table 1).

### UV mutagenesis and screening

One-week old protonemata from the *nog1dis* mutant was digested for 1 h in 1% Driselase dissolved in 8% mannitol. The resulting cell suspension was passed through a 40 µm cell strainer and centrifuged for 3 min at 120 xg without braking. Protoplasts were subsequently washed twice in 8% mannitol, with repeated centrifugation steps in between washes. Cells were resuspended in 6 ml 8% mannitol, counted using a haemocytometer, and then plated at a density of 50,000 cells per plate onto cellophane overlaid PRMB. Protoplasts were immediately exposed to a 75,000 mJ dose of UV light using a Stratalinker UV Crosslinker and this was performed with the petri dish lids removed. Following irradiation, the lids were quickly replaced, and the plates were wrapped with micropore tape. Plates were incubated at 24°C in the dark for 24 h, to prevent photoactivatable DNA damage repair, before being transferred to standard growth conditions for a further 2-3 weeks. When visible, regenerating protoplasts were transferred into individual wells of 24 well plates containing BCD medium. After approximately one month of growth in standard conditions, the mutagenized plants were screened for reversion of the *nog1dis* phenotype (i.e., restoration of gametophore development).

### Imaging

To stain cell walls, tissues were submerged in propidium iodide (10 µgmL^-1^) for around 10 min and then mounted on slides in water. Images were acquired using a Leica SP5 scanning confocal microscope with a 40x water immersive lens. A 488 nm laser was used to excite the propidium iodide with 30% laser power and fluorescence was detected at 600-630 nm.

### Plant phenotyping

To assay the number of gametophores produced by different *P. patens* lines, tissues were grown on cellophane-overlaid BCDAT plates for one week and then the tissues were harvested into separate 50 ml tubes containing 10 ml sterile water. The tissues were each homogenized with an IKA disperser for 20 s, and then 1 ml of homogenate was removed to measure optical density (OD600) in a spectrophotometer. The density of the homogenate was normalized by adding the required amount of water to each tube. 50 µl of homogenate was pipetted onto BCD medium in small petri dishes, with 5 repeats for each line. These plates were grown for two months under standard conditions, so that mature gametophores had sufficient time to develop. The number of gametophores formed on each plate were counted. To measure gametophore height, a representative and intact sample from each line was extracted from the tissue, placed in a petri dish and photographed. The heights were measured in ImageJ, by drawing a line from the apex to the base. To determine the cytokinin responsiveness of each line, tissues were grown on BCD medium for one week and then the cellophanes were cut and transferred to BCD medium containing 1 µM BAP for 3 days (or control medium) before being examined and images captured using a Leica M165C stereomicroscope equipped with a QImaging Micropublishing 5.1 RTV camera.

### Plant crossing and bulk segregant analysis

Both the *snog1a* mutant and the Reute::mCherry line (Perroud et al., 2019) were grown in magenta pots on BCD medium in standard growth conditions for 2-3 months. Pots were subsequently transferred to sporophyte induction conditions (16 h dark: 8 h light, 16°C). After approximately three weeks, gametophores were examined for the presence of gametangia (archegonia and antheridia). When gametangia were present, the Reute::mCherry line was submerged under water to make a sperm suspension, which was subsequently added to the *Ppsnog1a* mutant to allow outcrossing. After approximately 4-6 weeks, the resulting sporangia were removed carefully using forceps and examined under a Leica M165C fluorescence stereomicroscope to detect mCherry expression. This demonstrated a successful outcrossing event. Sporangia were sterilized in 70% ethanol for 4 min, washed 3 times with sterile water and then incubated at 4°C in the dark for one week. In a laminar flow hood, the sporangia were transferred to a 15 ml tube containing 10 ml of sterile water and then the sporangia ruptured with a pipette tip. 1 ml of spore suspension was plated onto each of ten cellophane-overlaid BCDAT plates and then grown in standard conditions to permit germination and subsequent growth. Individual sporelings were then transferred to wells of 24-well plates containing BCD medium and grown for a further 4-6 weeks. The progeny were screened for the presence or absence of gametophores, and sorted into two populations; *nog1dis*-like (wild-type pool, 80 individuals) and wild type or *snog1a*-like (mutant pool, 98 individuals). Genomic DNA was then extracted from one-week old protonemal tissues prepared from each individual line.

### Isolation of genomic DNA

Immediately before use, 0.07% (v/v) 2-mercaptoethanol and 0.1% (w/v) ascorbic acid was added to aliquots of extraction buffer (100 mM Tris-HCl pH 8.0, 1.42 M NaCl, 2 % CTAB, 20 mM EDTA, 2% PVP-40). The aliquots of extraction buffer were pre-warmed at 65°C in a water bath. One plate of 1-week-old protonemal tissues, grown on cellophane-overlaid BCDAT plates, was harvested and then blotted on filter paper to remove excess water. Tissues were then placed in a 2 ml microcentrifuge tube and frozen in liquid nitrogen. A miniature pestle was used to grind the tissue into powder, and then 500 µl of pre-warmed extraction buffer was added to the powder while frozen and further homogenised. An additional 200 µl of extraction buffer was added to the tube, along with 7 µl 10 mgmL^-1^ RNase A, and then incubated at 65°C for 10 min. 600 µl of chloroform-isoamyl alcohol (24:1) was added and the tube, shaken and centrifuged at 13,000 rpm for 10 min. Then the upper aqueous phase of the mixture was transferred to a fresh tube, 0.7 volumes of isopropanol were added, the tube shaken and spun again immediately at 13,000 rpm for 10 min. The pellet was washed in 70% ethanol, allowed to air-dry, and then resuspended in 30 µl nuclease-free water. A NanoDrop™ spectrophotometer was used to assess the quantity and quality of the extracted DNA.

### Preparation of genomic DNA samples for Whole Genome Sequencing

In total, four genomic DNA samples were prepared for sequencing; both parental lines (*nog1dis* and Reute::mCherry) and two pooled samples (a *snog1a*-like ‘mutant pool’ and a *nog1*-like ‘wild-type pool’). The pooled samples were prepared by pooling 1 µg genomic DNA extracted from all individuals in that population (80 individuals in the ‘WT pool’ and 98 individuals in the ‘mutant pool’).

### Candidate identification through bulk segregant analysis

Whole genome was performed using a NovoSeq X Plus sequencing platform (150 bp PE read lengths, 44X coverage) at Novogene. Bioinformatic analysis was performed as described previously (Moody et al., 2018; Moody et al., 2021). The presence of the premature termination codons in Pp3c8_19720 was confirmed by sequencing PCR products, amplified from both wild type and snog1a derived genomic DNA, using the primers Pp3c8_19720_int_F and Pp3c8_19720_int_R (Table 1).

### Physcomitrium patens transformation

Before the transformation, large quantities of plasmid DNA were acquired by midiprep using a QIAGEN plasmid midi kit. Approximately 20 µg of the plasmid was linearized by restriction digest overnight, treated with CIAP and precipitated with sodium acetate. Before starting the transformation, all solutions required were made fresh and filter sterilized using [filter]. The transformation was carried out in a biological safety cabinet. 2 g of polyethylene glycol (PEG) 6000 in a flat-bottomed vial, that had been sterilised in the autoclave, was melted in the microwave for approximately 1 min. 5 ml mannitol/Ca(NO_3_)_2_ solution (0.8% mannitol, 0.1 M Ca(NO_3_)_2_, 10 mM Tris pH 8.0) was added to the molten PEG 6000, shaken and allowed to cool for 2-3 h.

For each *P. patens* line that was to be transformed, 1-2 plates of tissue were harvested. 0.1 g Driselase enzyme was dissolved in 10 ml 8% mannitol for each digest. The tube containing the enzyme solution was wrapped in foil and rocked for approximately 10 min at room temperature. The tube was spun for 3 min in a centrifuge, and the supernatant was filter sterilised. *P. patens* tissue was placed in the tube with the Driselase solution, wrapped in foil and rocked very gently until the tissue appears well digested (at least 40 min). The digested tissue was put through a 70 µm cell strainer to separate protoplasts from remaining debris. The protoplasts were spun at no more than 120 xg, the supernatant was removed and the protoplasts were resuspended in 6 ml mannitol. This wash was repeated twice more. The cell density of the protoplasts was determined using a haemocytometer. The protoplasts were spun down again and resuspended in MMM (0.5 M mannitol, 0.15 M MgCl_2_, 0.1% MES pH5.6), to achieve a cell density of 1.5x10^6^ cells mL^-1^. The next steps were carried out without delay as the protoplasts cannot tolerate the MMM solution for very long. At least 10 µg of the linearised construct was pipetted into the bottom of a round-bottomed Falcon tube. To this was added 300 µl of protoplast suspension. 300 µl of the PEG solution was added in drops and the tube was swirled gently after the addition of each drop. The tube was heat-shocked at 45 °C in a water bath for 5 min and then incubated at room temperature for a further 5 min. Next, 300 µl 8% mannitol was added to the tube, 5 times at 3 min intervals and the mixture was gently swirled after each addition. Then, 1 ml 8% mannitol was added to the tube 5 times at 3 min intervals, gently tilting the tube after each addition to mix it. The tube was centrifuged at 120 xg, supernatant removed, and the cells were resuspended in 3 ml 8% mannitol. This suspension was gently pipetted onto three PRMB plates overlaid with cellophanes (1 ml for each plate). These plates were sealed with micropore tape and wrapped in foil for 24 h. Subsequently, they were unwrapped and placed in normal growing conditions for 5-7 days. Then the cellophanes, with the regenerating protoplasts, were transferred to selective BCDAT plates containing the appropriate selective agent. After another week on selective plates, the cellophanes are transferred to BCDAT plates with no selective agent to allow recovery. After another 1-2 weeks, the surviving *P. patens* colonies were individually transferred to selective BCD plates, to select for stably transformed colonies. Following another 1 – 2 weeks on selection, colonies that survived both rounds of selection were transferred to non-selective BCD plates and allowed to grow and generate tissue for further study.

### Generation of *snog1a* complementation lines

RNA was extracted form two-week-old wild-type tissue using an RNeasy kit (Qiagen) and treated with Turbo DNase (Ambion), according to the manufacturer’s specifications. cDNA was then synthesised using Superscript^TM^ III Reverse Transcriptase (ThermoFisher Scientific). Subsequently the Pp3c8_19720 transcript was PCR-amplified (excluding the stop codon) using Pp3c8_19720.FSalI and Pp3c8_19720.R_NOSTOP_HindIII and ligated into pZAG1 to create Act1p::Pp3c8_19720-GFP. This construct was linearised with SacII for subsequent transformation into the *snog1a* mutant. Stable transformants were selected using 100 µgml^-1^ Zeocin (Invitrogen; R25001). Genotyping was performed using the primers putativeSNOG1A_genomic2_F and putativeSNOG1A_genomic2_R (Table 1, Fig. S6).

### Generation of a *snog1a* deletion mutant

A deletion construct was designed. The construct consisted of the PpFLOE2L-1 5’ flanking sequence, with a small section of the PpFLOE2L-1 CDS and a PpFLOE2L-1 3’ flanking sequences. The 5’ and 3’ sequences were either side of a G418 resistance cassette. The construct was synthesized by TWIST Bioscience. The product was verified by Sanger sequencing. Prior to transformation into protoplasts isolated from the *nog1dis* mutant, the plasmid was linearized with PvuI. Stable transformants were selected using 40 µgml^-1^ G418. 5’ integration of the deletion construct was confirmed using the primers snog1a_del_genotyping_F and snog1a_del_genotyping_R (Table 1, Fig. S7).

### Phylogenetics

For each of the species included in the analysis, proteome sequences (primary transcript only) were obtained. The data used were: *Physcomitrium patens* v3.3 (Lang et al., 2018), *Marchantia polymorpha* v3.1 (Bowman et al., 2017), *Selaginella moellendorffii* v1.0 (Banks et al., 2011), *Oryza sativa* v7.0 (Ouyang et al., 2007), *Zea mays* PH207 v1.1 (Hirsch et al., 2016), *Sorghum bicolor* v3.1.1 (McCormick et al., 2018), *Brachypodium distachyon* v3.1 (Vogel et al., 2010), *Arabidopsis thaliana* Araport11 (Cheng et al., 2017), *Solanum lycopersicum* ITAG2.4 (Sato et al., 2012), *Medicago truncatula* Mt4.0v1 (Tang et al., 2014), *Populus trichocarpa* v3.1 (Tuskan et al., 2006), *Micromonas pusilla* CCMP1545 v3.0 (Worden et al., 2009) and *Ostreococcus lucimarinus* v2.0 (Palenik et al., 2007), *Chara braunii* S276v1.0 (Nishiyama et al., 2018), *Chlamydomonas reinhardtii* v5.5 (Merchant et al., 2007), *Amborella trichopoda* v1.0 (Albert *et* al., 2013), *Botryococcus braunii v2.1* (Browne *et al*., 2017), *Anthoceros agrestis* Oxford (Li *et al*., 2020), *Azolla filiculoides* v1.1 (Li, *et al*., 2018), *Brassica rapa* FPsc v1.3 (DOE-JGI, http://phytozome.jgi.doe.gov/), *Ceratodon purpureus* R40 v1.1 (Carey *et al*., 2021), *Ceratopteris richardii* v2.1 (Marchant *et* al., 2022), Chlorokybus atmophyticus CCAC 0220 v1.1 (Wang *et al*., 2020), *Klebsormidium nitens* NIES-2285 v1.1 (Hori *et al*., 2014), *Mesostigma viride* NIES-296 (Liang *et al*., 2020), *Spirogloea muscicola* CCAC 0214 (Cheng *et al.,* 2019), *Mesotaenium endlicherianum SAG 12.97* (Cheng *et al.,* 2019), *Coccomyxa subellipsoidea* C-169 v2.0 (Blanc *et al.,* 2012), *Dunaliella salina* v1.0 (Polle, *et al*., 2017), *Volvox carteri* v2.1 (Prochnik *eta l*., 2010), *Porphyra umbilicalis* v1.5 (Brawley *et al*., 2017), *Sphagnum fallax* v1.1 (Healey *et al*., 2023), *Diphasiastrum complanatum* v3.1 (DOE-JGI, http://phytozome-next.jgi.doe.gov/), *Gingko biloba* v2021 (Liu *et al*., 2021), *Glycine max* Wm82 ISU-01 v2.1 (DOE-JGI, http://phytozome.jgi.doe.gov), *Gossypium raimondii* v2.1 (Paterson *et al*., 2012), *Musa acuminata* v1 (D’Hont *et al*., 2012), *Panicum hallii* v3.2 (Lovell *et al*., 2018), *Salvinia cucullata* v1.2 (Li *et al*., 2018), *Spirodela polyrhiza* v2 (Wang *et al*., 2014), *Thuja plicata* v3.1 (Shalev *et al*., 2022), *Vitis vinifera* v2.1 (Jaillon *et al*., 2007), *Saccharomyces cerevisiae* R64-1-1 (Liachko et al., 2013), *Drosophila melanogaster* BDGP6.32 (Adams et al., 2000) and *Homo sapiens* GRCh38 (Lander et al., 2001).

With this set of proteomes, OrthoFinder was used to identify orthogroups (Emms & Kelly, 2019). The orthogroup containing *Pp3c8_19720* was selected and the protein sequences were aligned using MAFFT (L-IN-SI method) and a gene tree constructed using IQ-TREE, with automatic model selection (ModelFinder) and ultrafast bootstrapping (UFBoot) with 1000 replicates (Hoang et al., 2018; Kalyaanamoorthy et al., 2017; Katoh & Standley, 2013; Nguyen et al., 2015). Tree was rooted and edited in Interactive Tree of Life (iTOL) (Letunic & Bork, 2016).

## ACKNOWLEDGEMENTS

We are grateful to Pierre-François Perroud for proving the Renute::mCherry line, Yuji Hiwatahi for providing the pZAG1 plasmid; Andrew Cuming and Yasuko Kamisugi for providing the pAHG1 plasmid and John Baker for photography.

## AUTHOR CONTRIBUTIONS

Z.W. conducted the experiments, with assistance from G.C., E.D. and L.A.M.; L.A.M. conceived and designed the study; S.K. carried out the bioinformatics; Z.W. constructed the phylogenetic tree; and L.A.M. and Z.W. wrote and edited the manuscript.

## FIGURE LEGENDS

**Supplementary Figure 1. Generation of the *nog1dis* line.** A) Schematic of the construct designed to knockout the endogenous *PpNOG1* gene, and schematic of the modified *PpNOG1* locus in *nog1dis* following recombination. B) RT-PCR reveals the presence of the PpNOG1 transcript in wild type but not in the *nog1dis* mutant (tubulin – control).

**Supplementary Figure 2. Gametophores formed in the *snog1a* mutant are stunted relative to wild type.** Representative images of 2-month-old gametophores from wild type and *snog1a* plants. Scale bars, 0.5 cm.

**Supplementary Figure 3. Bulk segregant analysis and the identification of the causative mutation in the *snog1a* mutant.** A) An outcrossing event between snog1a and the Reute::mCherry line yields a diploid sporophyte that undergoes meiosis to produce phenotypically segregating progeny (phenotypic outcomes highlighted). (B,C) Expected snog1a mutant, SNOG1A WT, nog1 mutant and NOG1 WT allele frequencies in the mutant (B) and wild-type (C) pools respectively. D) Allele frequency plot for segregants on chromosome 8 of the *P. patens* genome assembly.

**Supplementary Figure 4. Phylogenetic analysis of FLOE-related homologues in the green lineage.** Bootstrap values have been indicated on each branch. Both FLOE1L and FLOE2L clades have also been indicated.

**Supplementary Figure 5. Alignment of AtFLOE1, AtFLOE2, AtFLOE3 and PpFLOE2L-1.** Conserved domains have been highlighted as indicated.

**Supplementary Figure 6. Generation of the *snog1a* complementation line.** A) Schematic of the construct used to complement the *snog1a* mutant phenotype, and the resulting targeted locus. B) Genotyping of the complementation line using putativeSNOG1A_genomic2_F and putativeSNOG1A_genomic2_R primers denoted by purple and green arrows in (A) respectively. The construct is only detected in the complemented line and not in wild type (tubulin – control).

**Supplementary Figure 7.** (A) Schematic of the construct used to disrupt the *PpFLOE2L-1* locus in the *nog1dis* mutant, and the resulting targeted locus. B) Genotyping of the complementation line using snog1a_del_genotyping_F and snog1a_del_genotyping_R primers denoted by blue and red arrows in (A) respectively.

